# DeepPPPred: An Ensemble of BERT, CNN, and RNN for Classifying Co-mentions of Proteins and Phenotypes

**DOI:** 10.1101/2020.09.18.304329

**Authors:** Morteza Pourreza Shahri, Katrina Lyon, Julia Schearer, Indika Kahanda

## Abstract

The biomedical literature provides an extensive source of information in the form of unstructured text. One of the most important types of information hidden in biomedical literature is the relationships between human proteins and their phenotypes, which, due to the exponential growth of publications, can remain hidden. This provides a range of opportunities for the development of computational methods to extract the biomedical relationships from the unstructured text. In our previous work, we developed a supervised machine learning approach, called PPPred, for classifying the validity of a given sentence-level human protein-phenotype co-mention. In this work, we propose DeepPPPred, an ensemble classifier composed of PPPred and three deep neural network models: RNN, CNN, and BERT. Using an expanded gold-standard co-mention dataset, we demonstrate that the proposed ensemble method significantly outperforms its constituent components and provides a new state-of-the-art performance on classifying the co-mentions of human proteins and phenotype terms.

---

Proteins are considered one of the most important biomolecules, which are critical for the maintenance and development of life [3]. A cell’s full set of expressed proteins–the proteome–is dynamic and multidimensional with these proteins operating in a complex network and ensures the integrity of cellular structure and function [17]. Errors in the underlying genetic sequence of the protein often cause alterations in critical regions of a protein’s structure. Such errors can alter the protein’s function-specific tertiary structure, which results in changes to its phenotypes [10]. In the medical context, a phenotype is defined as a deviation from normal physiology or behavior [35]. Well-known changes in phenotype are brought about by alterations in structure or regulation of one or more proteins involved in important biological pathways, such as those relevant to Alzheimer’s disease, cancer, cystic fibrosis, Huntington’s disease, and type II diabetes, Parkinson’s disease [3, 11, 23]. Uncovering novel changes in protein structure, function, and regulation–in addition to understanding how these alterations lead to human disorders–is a popular field of research in the biomedical community [3, 5, 11, 23, 38, 34, 10, 17].

Human Phenotype Ontology (HPO) is a standardized vocabulary that covers a wide range of phenotypic abnormalities associated with human diseases [14]. HPO contains several sub-ontologies, and its main sub-ontology is *Phenotypic abnormalities* that represents clinical abnormalities. Each sub-ontology includes HPO terms and associated HPO identifiers (IDs), e.g. *Parkinsonism*, HP:0001300. Each sub-ontology has a hierarchical structure where more general terms appear at the top, and more specific terms are closer to the leaves. Each pair of terms in the hierarchy are connected with a *is-a* relationship. In this paper, we use *phenotypes* and *HPO terms* interchangeably. HPO website^1^ provides gold-standard annotations for a large collection of human proteins acquired through biocuration. Biocuration is the process of extracting knowledge from unstructured text and storing them in knowledge bases. It is usually performed manually with the help of computational tools [1]. However, currently, only a small portion of known human proteins have HPO annotations [14]. Nevertheless, researchers believe that many other human proteins are associated with diseases and should be annotated with HPO terms (Peter Robinson, personal communication, 2015).

Expanding knowledge bases such as the HPO database through biocuration is essential for potential future applications in medicine and healthcare. However, biocuration is considered slow and resource-consuming. Thus, to facilitate the typically slower rate of human annotation, efficient and accurate computational tools are required to expedite this process [1]. Consequently, researchers who work in the area of biomedical relationship extraction have shown interest in developing computational models to extract relationships between proteins and phenotypes [40, 15, 9, 13].

As a solution to the above, in a recent study [31], we proposed a novel two-step approach for extracting human protein-HPO term relationships. The first step was to extract protein-HPO *co-mentions*, which are co-occurrences of protein names and phenotype names in a particular span of text, i.e., a sentence, a paragraph, etc [28]. In our previous work, we developed ProPheno^2^, which is an online and publicly accessible dataset comprising proteins, phenotypes (HPO terms), and their co-occurrences (co-mentions) in text. These co-mentions are extracted from Medline abstracts and PubMed Central (PMC) Open Access full-text articles by using a sophisticated knowledge-free Natural Language Processing (NLP) pipeline [30]. This dataset covers all terms in the *Phenotypic abnormality* sub-ontology. However, a knowledge-free Natural Language Processing pipeline extracts every co-mention of proteins and phenotypes, but not all protein-phenotype co-mentions imply that there is a relationship between the two entities (see Fig. 1 for an example).

**Fig. 1.**
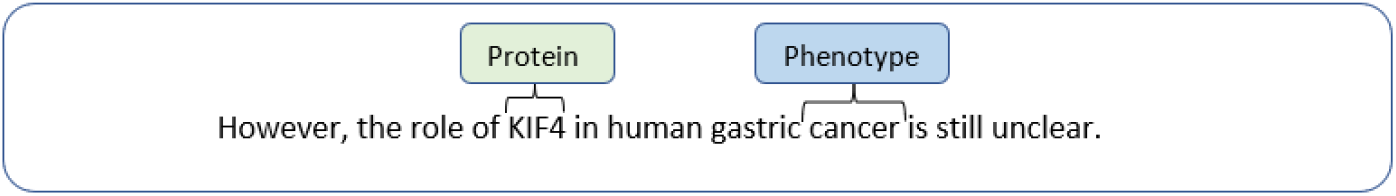
An example of a *bad* co-mention in which the sentence does not convey a relationship between the protein, i.e. “KIF4”, and the phenotype, i.e. “cancer”. (PMID: 20711700)

Therefore, in the second step of this two-step approach, extracted co-mentions are filtered using a co-mention classifier that can distinguish between good and bad co-mentions. We define a co-mention as a *good* co-mention if there is enough evidence mentioned in the text indicating a relationship between the protein and the phenotype. In other words, a good co-mention is a valid relationship between the two entities based on the meaning of the context text. Fig. 2 depicts an example of a good co-mention of a protein and a phenotype in a sentence. The combination of a co-mention extractor and a co-mention classifier/ filter constitutes a complete relationship extraction pipeline.

**Fig. 2.**
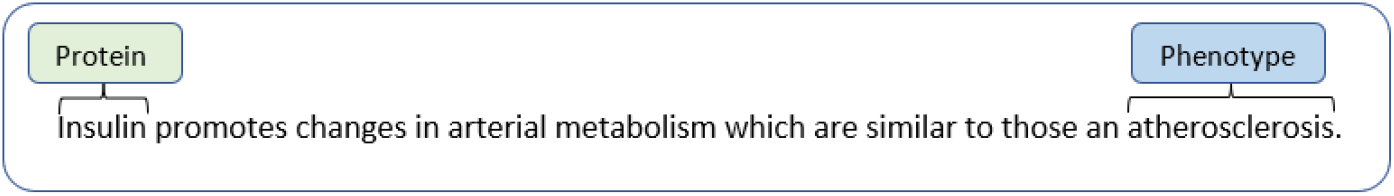
An example of a good sentence-level protein-phenotype co-mention, which is extracted from the article PMID: 18596936.

In the same study, [31], we developed PPPred (Protein-Phenotype Predictor), a co-mention classifier for classifying protein-phenotype co-mentions. We first randomly selected a subset of co-mentions from the ProPheno database and curated it with the assistance of biologists. This gold-standard dataset was composed of 809 human protein-HPO term co-mentions annotated with binary labels of good/ bad. Then, we used this dataset for developing predictive models using traditional supervised machine learning techniques. PPPred was based on Support Vector Machines (SVMs) and employed an extensive collection of syntactic and semantic features. While PPPred significantly out-performed other baseline methods according to our experimental results, there was significant room for improvement.

In our current work presented in this paper, we first double the gold-standard dataset by curating an additional set of 800+ co-mentions with the assistance of two biologists. This allows us to apply data-hungry deep learning techniques for the task of protein-phenotype co-mention classification. Next, we develop three such deep learning models: a BERT (Bidirectional Encoder Representations from) [8] model adapted for our tasks, and a CNN and an RNN model specifically designed for this task. Finally, we create an ensemble model composed of these three deep learning models and PPPred. Ensemble learning has several advantages: 1) They generally provide higher performance than their components. 2) Model selection for a problem can be tedious, but ensemble methods combine several models so that the model selection may be circumvented. 3) Learning a single model on very large data can be difficult [27].

Our experimental results suggest that the proposed ensemble model can outperform all its constituent deep learning models as well as PPPred. The expanded dataset is also made publicly available^3^ [29] for the benefit of the community. Additionally, we made available all our codes publicly available^4^. To the best of our knowledge, this is the first study of developing deep learning models for human protein-HPO term co-mention classification.

The rest of the paper is organized as follows. Section 1 provides a brief background on the related work in this area. The proposed method is discussed in Section 2. Section 3 discusses the results of running this method and compares the results with other methods and provides a discussion on the results. Finally, Section 4 concludes the study and discusses future work and open problems.

## 1 Related Work

The main approaches for biomedical relationship extraction include co-occurrence-based methods, rule-based methods, and machine learning-based methods. Co-occurrence methods look for any co-mention of the two entities of interest in a particular span of text, e.g., sentence, paragraph, etc., and usually provide low precision and high recall values [4]. Rule-based methods define linguistic patterns and extract the relationships using the patterns [2, 21, 33, 24, 12]. The rules can be derived from manually annotated corpora using machine learning algorithms or defined manually by a domain expert. Several studies focus on employing lexical analyzers and parsers to identify the relationships between entities [37, 41, 44, 7].

Machine learning-based approaches are also employed for the relationship extraction from biomedical text [39, 16, 13, 20]. The machine learning category includes methods based on feature engineering, graph kernels, and deep learning. Support Vector Machines (SVMs) have shown high performance in biomedical relationship extraction, but they need feature engineering, which is a skill-dependent task [45]. Kernel-based methods also require designing suitable kernel functions. Deep neural network-based methods eliminate the need for feature extraction and defining rules, and provide state-of-the-art on various tasks in biomedical relationship extraction [45, 26].

Deep neural networks have been widely used in biomedical relationship extraction. For example, Liu et al. [19] classify drug-drug interaction (DDIs) using CNNs with the help of word sequences and position sequences. Peng and Lu [25] utilize word sequences, positions sequences, POS tags sequences, and dependency vectors to classify protein-protein interatcions (PPIs) using two-channel CNNs. Quan et al. [32] propose a multichannel CNN for extracting various biomedical relationships. Lim and Kang [18] propose a tree-LSTM model for chemical-protien interactions (CPIs) classification. Wang et al. [42] present a dependency-based Bi-LSTM to classify DDIs. Corbett et al. [6] also classify CPI relationships using LSTMs. Besides, Sahu and Anand [36] combine Bi-LSTM with attention pooling to improve performance.

Furthermore, some researchers have combined RNNs and CNNs to create hybrid models. Peng et al. [26] introduce an ensemble of SVMs, CNNs, and RNNs, for the task of chemical-protein relationship extraction in BioCreative VI. The majority voting and Stacking schemes are used for combining the outputs of the three methods. While deep neural networks often show top performance on biomedical relationship extraction tasks (e.g. [26]), the main issue with deep neural networks is that they require large labeled datasets to provide superior performance.

Despite a large number of studies conducted on extracting entity relationships from the biomedical literature (including a handful of methods for extracting relationships between genes/proteins and phenotypes), the only method designed explicitly for human protein-HPO term relationship extraction is PPPred, which is a traditional machine learning classifier that was previously developed by our lab [31].

## 2. Methodology

### 2.1 Approach

Aligning with our previous work [31], we formulate the task of co-mention classification as a supervised learning problem as described below.

Given a context *C* = *w*_1_*w*_2_..*e*_1_..*w*_3_..*e*_2_..*w*_*n-*1_*w*_*n*_ composed of words *w*_*i*_ and the two entities *e*_1_ and *e*_2_, we define a mapping *f*_*R*_(·) as:

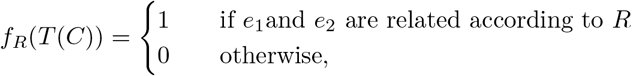

where *T*(*C*) is a high-level feature representation of the context, *e*_1_ and *e*_2_ are the entities representing the protein and the phenotype, and *R* is the relationship that represents the protein-phenotype relationship between the two. An example is considered a positive example if the meaning of the context suggests that the protein mentioned has this function (i.e., a good co-mention). Otherwise, it is labeled as a negative example.

In this work, the context *C* is a single sentence (i.e., the sentence containing the mentions of the two entities). Fig. 2 depicts a sentence which is labeled as a positive example (i.e., *f*_*R*_ = 1) because it provides evidence for the relationship between the two entities “JARID1C” (protein) and “aggression” (phenotype). We model this problem as a supervised learning problem and use binary classifiers for learning *f*_*R*_.

### 2.2 Data

In our previous work [31], we used ProPheno 1.0 [30], which is a dataset of proteins-phenotype co-mentions extracted from the biomedical literature. This dataset maps the proteins and phenotypes to the corresponding UniProt_5_ IDs and HPO IDs. In that work, we randomly selected a dataset of 809 sentence-level co-mentions of proteins and phenotypes from ProPheno. This dataset was then annotated by two biologists to generate the gold-standard dataset. The annotators were provided instructions to label a co-mention as good/ positive if the sentence conveys that the protein and the phenotype has a relationship. Otherwise, the co-mention was labeled bad/negative.

However, we observed that the above dataset was under-representative of the problem [31]. Nonetheless, we have access to millions of co-mentions through ProPheno [30]. Therefore, we annotated another random subset of co-mentions from ProPheno consisting of 876 sentences. For the new dataset, we did not include the most frequent proteins and phenotypes, which are “Neoplasm” (HP:0002664) and “Receptor tyrosine-protein kinase erbB-2”. The new subset was also annotated by two biologists, and we combined the two subsets, created a list of 1,685 annotated sentence-level co-mentions, and used this combined dataset as the gold-standard data for this work. Subsequently, we divided the list of sentences into three datasets, including training, validation, and test sets. The training set is composed of 1,010 (60%) sentences, the validation set contains 337 (20%) sentences, and the test set comprises 337 (20%) sentence-level co-mentions. The inter-annotator agreement is calculated using Cohen’s Kappa statistic [22], and the corresponding value is 0.64, which shows substantial agreement.

Tables 1, 2, and 3 demonstrate the distribution of classes in the training, validation, and test sets, respectively. According to the Table 1, 27% of sentences in the training set are extracted from the abstracts, and 73% are from the full-text articles. Among the sentences from the abstracts, 57% are labeled as “good” and 43% are labeled as “bad”. The distribution for the sentences from the full-text articles is 70% and 30% “good” vs. “bad”, respectively. The overall class distribution is 67% and 33% for “good” and “bad”, respectively. and phenotypes in the sentences. These plots also state that Good and Bad co-mentions are distributed similarly in the dataset.

**Table 1.** The class distribution in the training set

|  | Good | Bad | Total |
| --- | --- | --- | --- |
| Sentences from abstracts | 115 (57%) | 118 (43%) | 273 (27%) |
| Sentences from full-texts | 519 (70%) | 218 (30%) | 737 (73%) |
| All sentences | 674 (67%) | 336 (33%) | 1,010 |

**Table 2.**
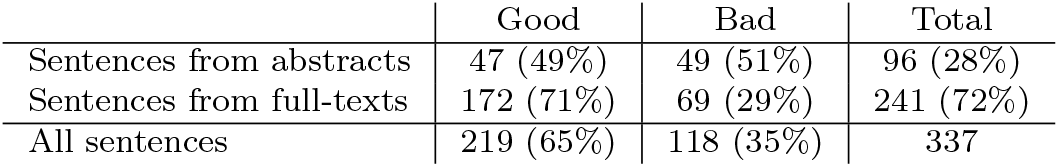
The class distribution in the validation set

|  | Good | Bad | Total |
| --- | --- | --- | --- |
| Sentences from abstracts | 47 (49%) | 49 (51%) | 96 (28%) |
| Sentences from full-texts | 172 (71%) | 69 (29%) | 241 (72%) |
| All sentences | 219 (65%) | 118 (35%) | 337 |

**Table 3.**
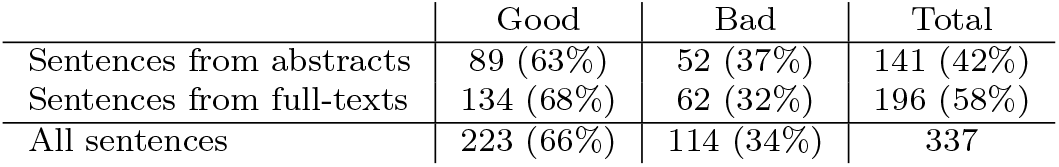
The class distribution in the test set

|  | Good | Bad | Total |
| --- | --- | --- | --- |
| Sentences from abstracts | 89 (63%) | 52 (37%) | 141 (42%) |
| Sentences from full-texts | 134 (68%) | 62 (32%) | 196 (58%) |
| All sentences | 223 (66%) | 114 (34%) | 337 |

Fig 3. depicts the distribution of the depths of HPO terms in the annotated co-mentions. The plots show that Good co-mentions and Bad co-mentions have similar distributions for various depth levels. Fig. 4 also provides the distribution of the lengths of the shortest dependency path between the proteins

**Fig. 3.**
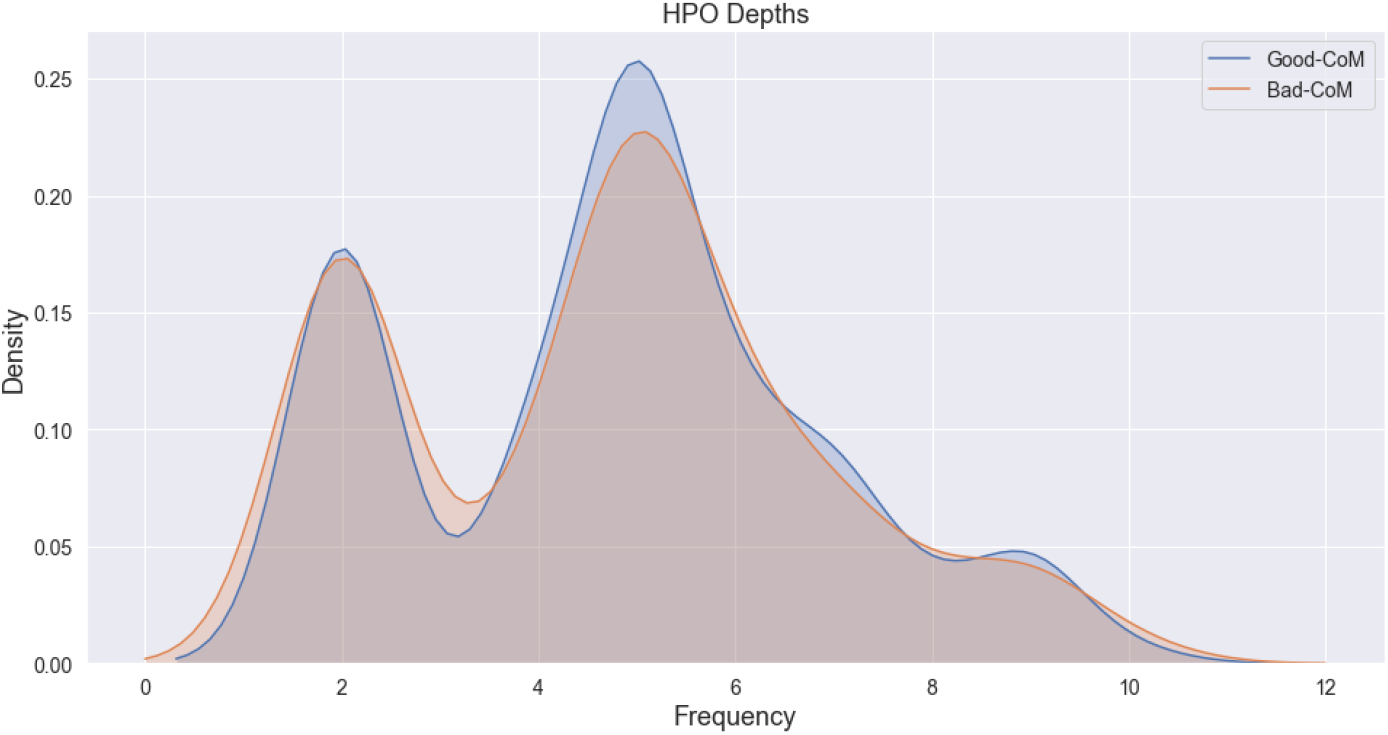
Distribution of depth of HPO terms in the annotated co-mention data

**Fig. 4.**
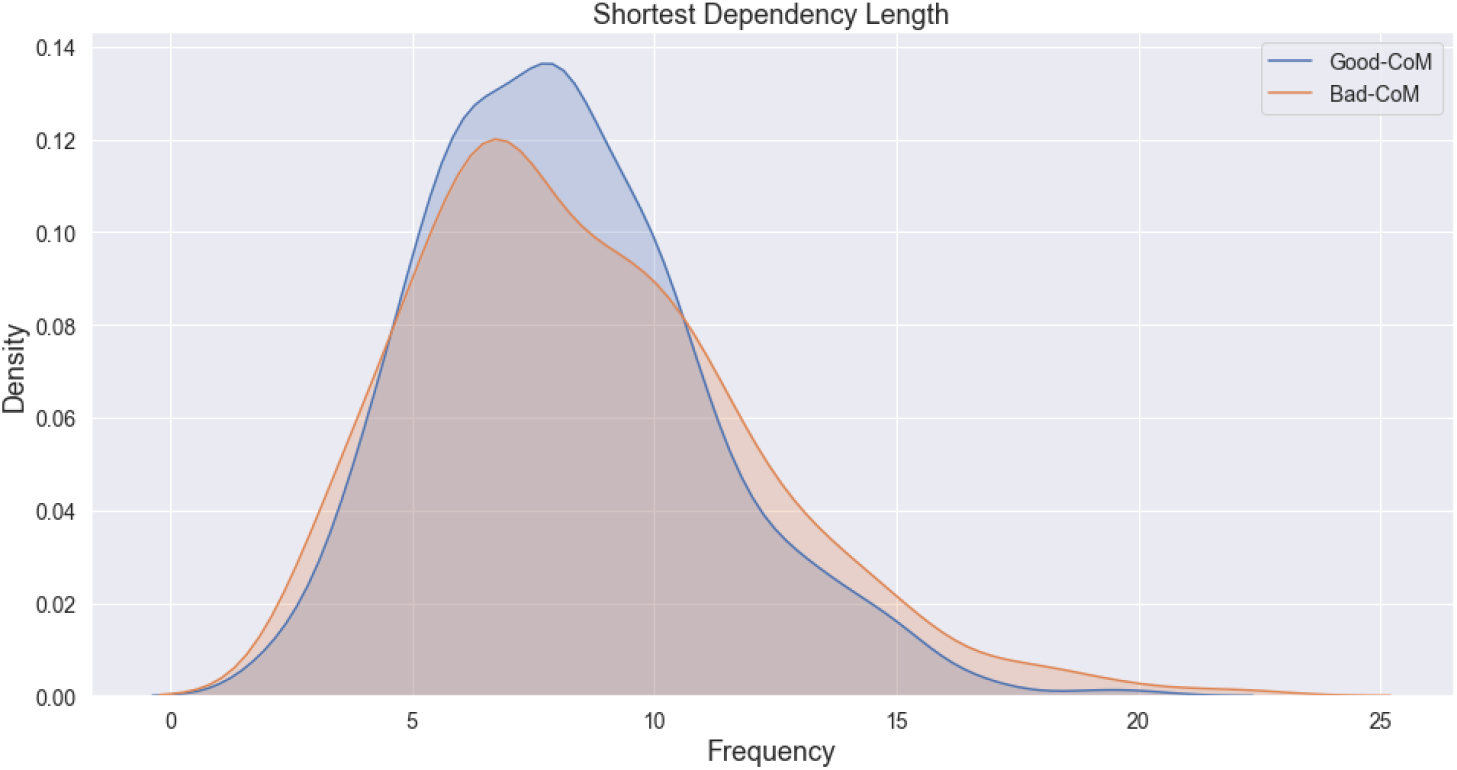
Distribution of length of shortest dependency path between the proteins and phenotypes in the annotated co-mention data.

Tables 4 and 5 show the most frequent phenotypes and proteins in the dataset, respectively. According to the tables, 25% of the sentences discuss the HPO term “Neoplasm” (HP:0002664) (other names: “Cancer” or “Tumour”), and 9% of the sentences mention the protein “Receptor tyrosine-protein kinase erbB-2” (P04626). Table 6 also demonstrates the most frequent protein-phenotype pairs mentioned in the dataset. We observe that 6% of the co-mentions in the dataset mention above protein-phenotype pair, which shows this pair is a well-studied protein-phenotype pair.

**Table 4.** Most frequent HPO terms mentioned in the dataset

|  | HPO ID | HPO Term | Depth | Frequency |
| --- | --- | --- | --- | --- |
| 1 | HP:0002664 | Neoplasm | 2 | 424 |
| 2 | HP:0003002 | Breast carcinoma | 5 | 138 |
| 3 | HP:0001909 | Leukemia | 4 | 78 |
| 4 | HP:0002861 | Melanoma | 4 | 51 |
| 5 | HP:0000819 | Diabetes mellitus | 6 | 27 |

**Table 5.** Most frequent proteins mentioned in the dataset

|  | UniProt ID | Protein Name | Freq. |
| --- | --- | --- | --- |
| 1 | P04626 | Receptor tyrosine-protein kinase erbB-2 | 149 |
| 2 | P01308 | Insulin | 66 |
| 3 | Q9Y617 | Phosphoserine aminotransferase | 57 |
| 4 | O14788 | Tumor necrosis factor ligand superfamily member 11 | 56 |
| 5 | P09486 | SPARC | 35 |

**Table 6.** Most frequent protein-HPO term pairs mentioned in the dataset

|  | UniProt ID | HPO ID | Protein Name | HPO Term | Frequency |
| --- | --- | --- | --- | --- | --- |
| 1 | P04626 | HP:0002664 | Receptor tyrosine-protein kinase erbB-2 | Neoplasm | 102 |
| 2 | Q9Y617 | HP:0002664 | Phosphoserine aminotransferase | Neoplasm | 54 |
| 3 | P04626 | HP:0003002 | Receptor tyrosine-protein kinase erbB-2 | Breast carcinoma | 37 |
| 4 | Q03164 | HP:0001909 | Histone-lysine N-methyltransferase 2A | Leukemia | 21 |
| 5 | O14788 | HP:0003002 | Tumor necrosis factor ligand superfamily member 11 | Breast carcinoma | 14 |

### 2.3 Preprocessing

The next step is to preprocess the data to make it ready for our models. Preprocessing includes replacing protein and phenotype entities, tokenization, converting the data into a numeric format, and padding/ truncating. The following sections discuss each step in more detail.

#### 2.3.1 Proteins and Phenotype Tokens

Some protein names and phenotype names are composed of multiple words that make interpretations difficult. We replace protein names and phenotype names in sentences with “PROT” and “PHENO”, respectively, to have a unique format across all sentences.

#### 2.3.2 Tokenizer

In the next step, we need to convert the text into a list of tokens. To obtain a tokenizer, we use Keras^6^ tokenizer that is capable of fitting a tokenizer on the unstructured text and converting the list of texts into a list of sequences. These sequences can be used as direct input to our model.

#### 2.3.3 Padding and Truncating

The sequences need to have the same shape to be used as the input to our model. Therefore, we need to truncate the sequences longer than a cutoff, and also, we need to add zeros to the end of sequences shorter than that specific value. We set the maximum length of sequences to a cutoff of 80, and we perform padding and truncating to make all of them the same size.

### 2.4 Models

Fig. 5 depicts the overview of the proposed deep ensemble model, which is capable of classifying sentence-level co-mentions of proteins and phenotypes from biomedical literature. In this model, inspired by a Peng et al.’s study [25], we start with training four separate classifiers on the training set. Subsequently, we make predictions using the trained models on the validation set. By feeding the predicted probabilities made by the models, we train an additional model using Logistic Regression-based *Stacking* [43] to combine the outputs of previous models. The stacking method learns how to combine the predictions of multiple models into one ensemble model.

**Fig. 5.**
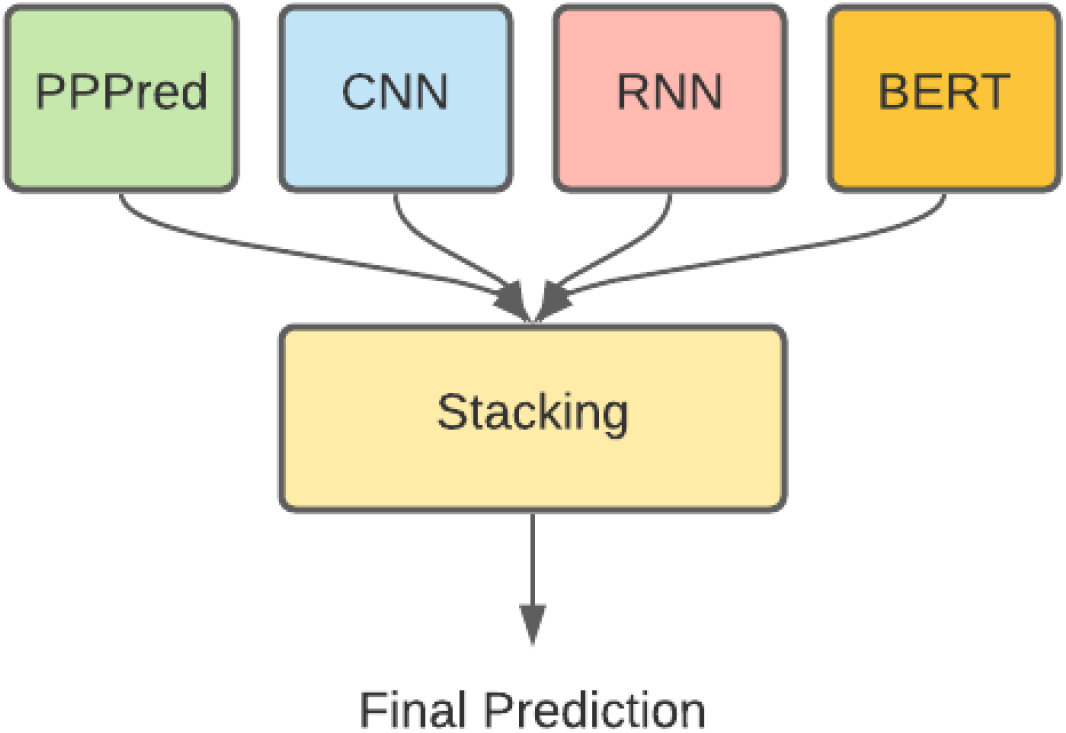
The proposed ensemble model that combines PPPred with BERT, a convolutional neural network, and a recurrent neural network.

The four models used for developing our ensemble classifier are: (a) PPPred [31] that we have developed previously, (b) a model that utilizes the BERT language model, which we fine-tune for this particular task, (c) a CNN model, and (d) an RNN model. Both the CNN and RNN models are specifically designed for this specific problem. The following sections describe the BERT, CNN, RNN models in detail.

#### 2.4.1 BERT Model

As mentioned earlier, BERT is already pre-trained on millions of articles, and it can be utilized for a variety of tasks. In this work, we fine-tune BERT for text classification by performing tokenization on sentences and adding “[CLS]” and “[SEP]” tokens to the start and end of the sequence. We fill the rest of the input sequence with zeros to make all the sequences the same length. BERT also requires the positional embeddings and segment embeddings as input that we pass them with the current sentence to BERT. After fine-tuning, we take the prediction from the output layer of BERT. Fig. 6 demonstrates the described BERT model. In this figure, the segment embeddings are filled with ones for all of the tokens since we pass the entire sentence as one input.

**Fig. 6.**
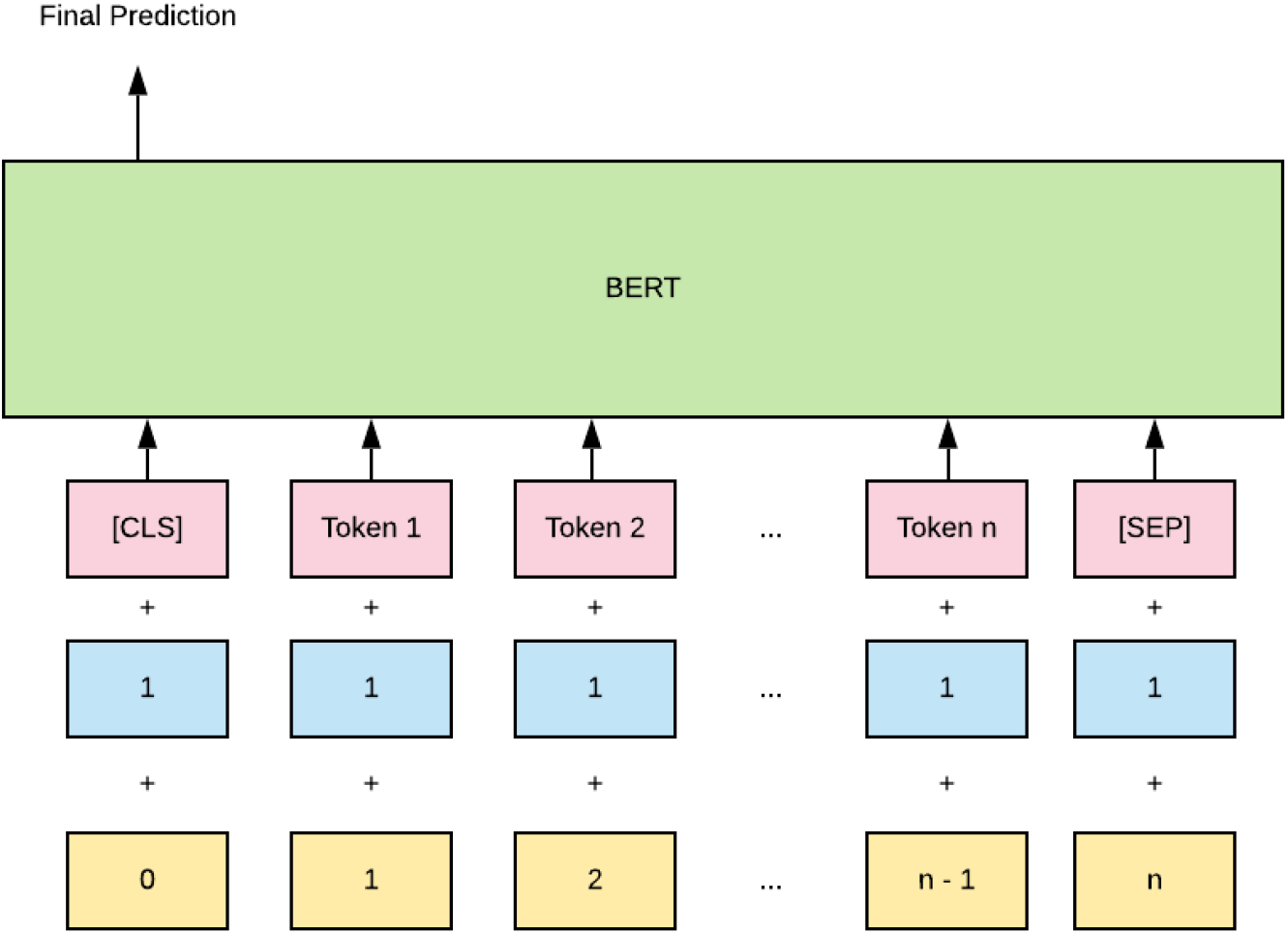
The proposed BERT model that takes the tokenized sentence, positional embeddings, and segment embeddings as input, and returns the probability of belonging the input to each class at the output layer.

#### 2.4.2 CNN Model

We propose a convolutional neural network inspired by Peng and Lu’s study [25]. We create 64 kernels for each network with window sizes of three and five. To prevent the model from overfitting, we employ dropout values of 0.3 and 0.5 for the original and shortest path sequences, respectively. After max pooling with a stride value of two, we flatten the sequences and concatenate them together. Next, we add two fully-connected layers with sizes of 100 and 64. For each fully-connected layer, we add a dropout layer with a value of 0.2. We use ReLU activation functions for all of the layers except the output layer in which we utilize the Sigmoid function to get the confidence score between zero and one. Fig. 7 demonstrates the described convolutional neural model.

**Fig. 7.**
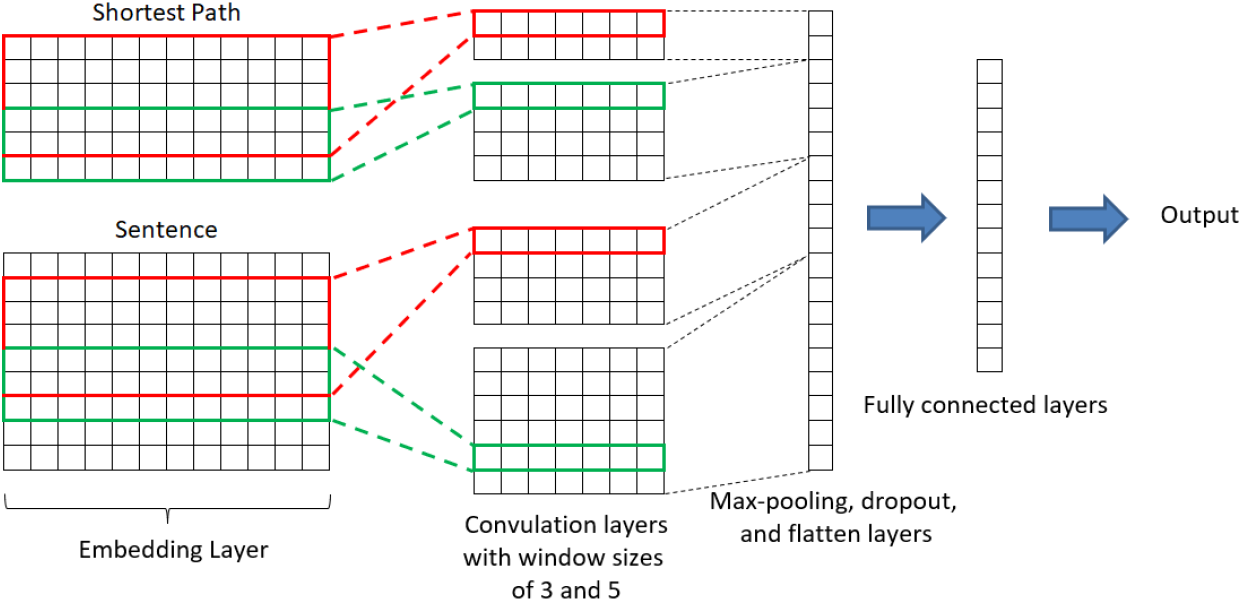
The proposed Convolution neural network

#### 2.4.3 RNN Model

We design another neural network for this specific task that is based on recurrent neural network architecture. We employ two Bi-directional LSTM layers for the original sequence and shortest path sequence with the sizes of 32 for each layer. Eventually, we add two fully-connected layers with the sizes of 100 and 64, and two dropout layers with a value of 20%. We pass the fully-connected layers through the ReLU activation function, and the Sigmoid function is used for the output layer. Fig.8 shows the described BiLSTM model.

### 2.5 Experimental Setup

We implemented all models using the PyTorch package^7^ and the SciKit-learn^8^ package. All the neural networks have 20 epochs and use binary cross-entropy loss function and the Adam optimizer. We also fine-tine BERT for four epochs. Eventually, we combine the predictions of four models using the Logistic Regression classifier on the validation set.

We perform 10-times repeated train/test validation with various seeds and average across them for comparing the performance of the proposed models with others. We report precision, recall, and F1 values as the primary performance measures. We also measure the significance of the difference in performance using paired t-tests.

The RNN model takes 240 seconds for training, while the CNN model can be trained in 90 seconds. The BERT model also takes 600 seconds to be fine-tuned on the training data. The average time for combining the outputs of these models using Stacking is two minutes.

## 3 Results and Discussion

Fig. 9 shows the comparison of the results of running the PPPred, RNN, CNN, BERT, and the Ensemble model on the test set. We observe that PPPred obtains the best precision value compared with other models, which can be due to the specific features we designed for this problem. However, the highest recall value, among the individual components of our ensemble model, is obtained by the BERT model. The BERT model also provides the best F1 score among the individual models. The RNN and BERT models achieve very close F1 scores, and it shows that RNNs, which are designed for sequence problems, perform similar to the BERT model, which provides state-of-the-art in several NLP tasks.

**Fig. 8.**
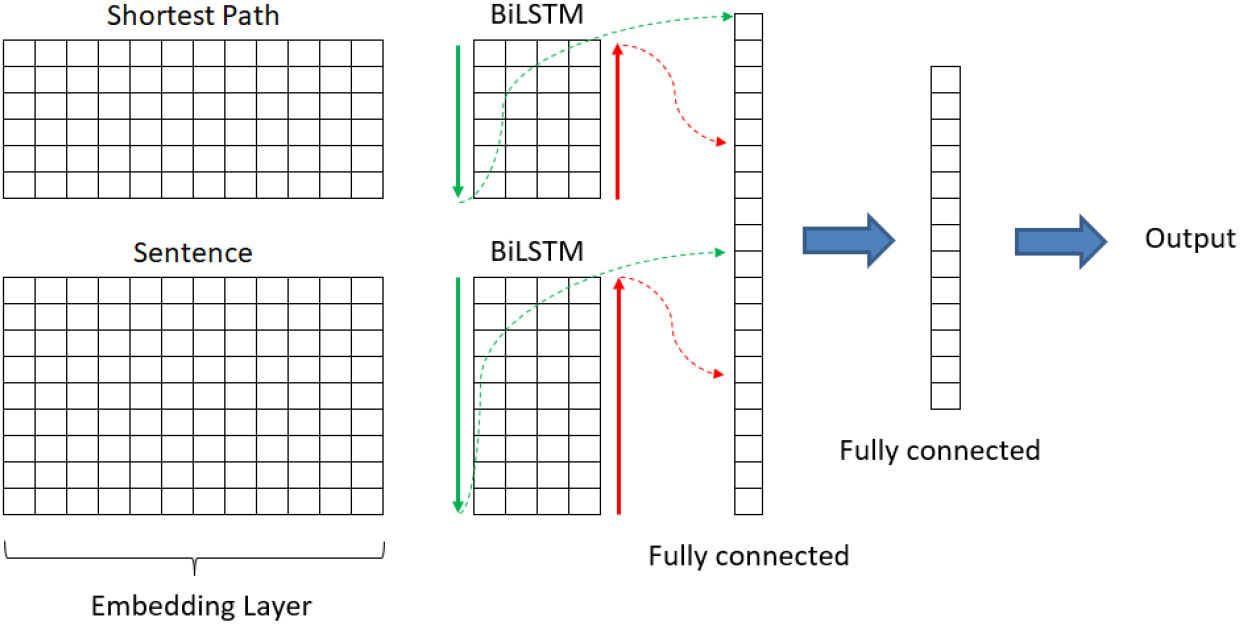
The proposed BiLSTM neural network

**Fig. 9.**
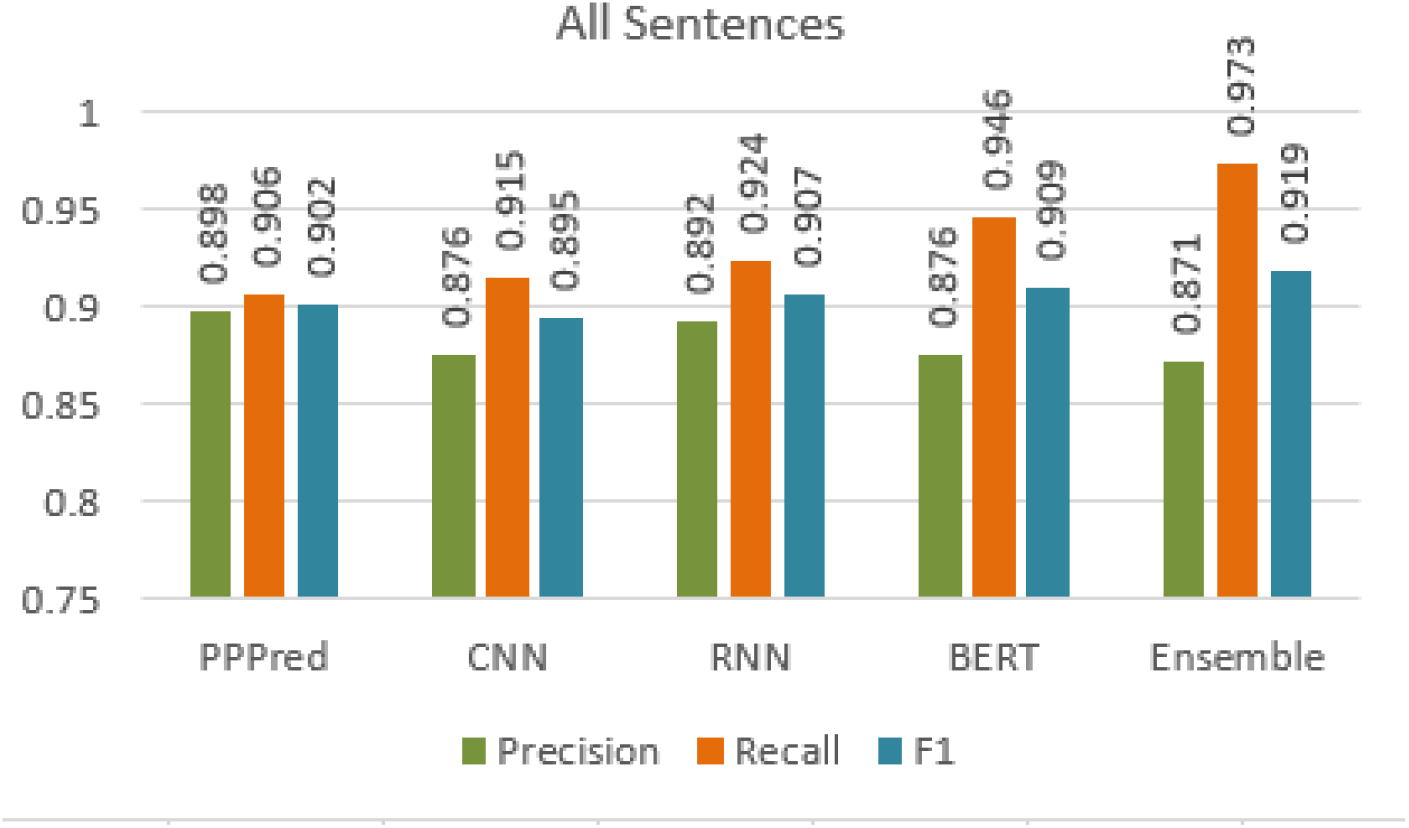
The comparison of multiple models’ performances on the test set.

Overall, we observe that the proposed ensemble model achieves the best F1 score (0.919) among all models. We also observe that the ensemble model significantly outperforms its components in recall as well. PPPred performs better than the CNN model, which automatically performs feature extraction.

It also obtains a comparable F1 score to the RNN model. This observation highlights the fact that engineered features for PPPred can provide as good performance as the new neural networks in the context of smaller datasets. However, a larger annotated dataset, which is hard to come by, may have tipped the scales in favor of CNN and RNN models.

We also investigate the separate performance of the described models on abstracts and full-text articles. Fig. 10 depicts the comparison of F1 scores obtained by the PPPred, CNN, RNN, BERT, and ensemble models on the sentences extracted from abstracts, full-text articles, and both. We observe that the proposed ensemble model provides the best F1 score on the sentences from the abstracts (0.902), whereas the PPPred model outperforms other models on the full-texts’ sentences. This is because of the high precision value obtained by the PPPred compared to other models.

**Fig. 10.**
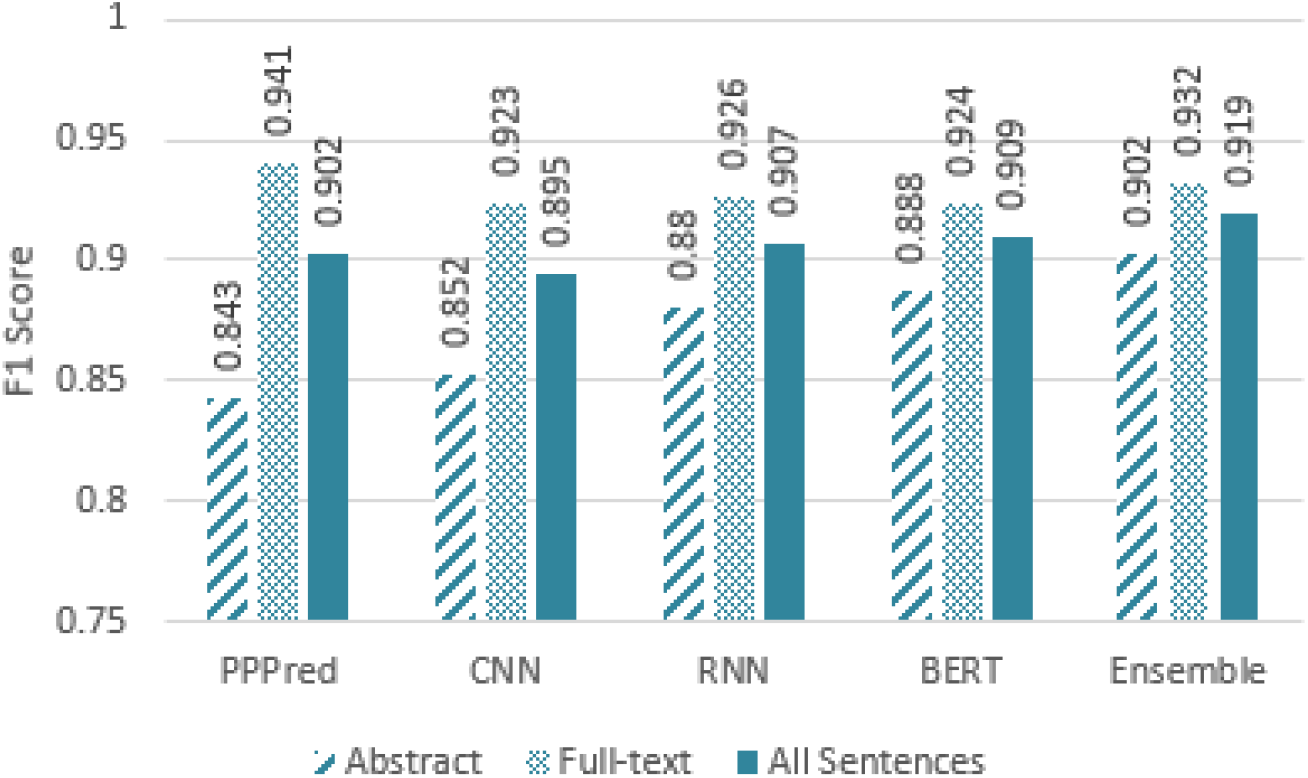
Comparison of F1 scores on the sentences from abstracts, full-text articles, and both.

Our results (data not shown) suggest that the CNN model performs better on shorter sentences, whereas the RNN model provides better performance on longer sentences. We observe that by using RNNs, the average length of wrong predictions is 231 characters, whereas the average length of wrong predictions made by the CNN model is 251 characters. The mean length of sentences is 219.8, and the standard deviation is 96.5. This is an intuitive observation since CNNs are designed to extract local features from the input. On the other hand, the added cell-state in LSTMs provides the ability to learn from long sequences. Peng et al. has observed the same [26].

Another observation from the results is that longer sentences are harder to classify (data not shown). Our designed models provide false positives and false negatives when the input sentence is long and contains multiple clauses and relationships between several bio-entities. For example, consider the following sentence: “It was on the basis of these concerns, and with a view to shedding further light on the expression of Reelin in human liver injury, that we investigated hepatic **Reelin** expression in a large series of patients with HCV-related chronic **hepatitis**, and verified its relationship with the other histological and immunohistochemical markers used to reflect activity and severity of liver disease.”. This is a long sentence that contains multiple phenotypes, and our ensemble model is unable to make the correct prediction.

## 4 Conclusions and Future Work

In this study, we created a co-mention classifier/filter that distinguishes between good and bad co-mentions of proteins and phenotypes in sentences. Specifically, we created an ensemble model composed of several supervised classifiers, including several sophisticated deep learning models, trained using manually-annotated sentence-level co-mentions of proteins and phenotypes. This ensemble classifier can be employed to perform highly accurate co-mention classification on protein and phenotype entities mentioned in biomedical literature. We observed that the proposed ensemble model provides the best performance compared with its individual classic and modern supervised components. We combined the BERT model that was a break-through in NLP, with other deep learning models and obtained state-of-the-art for the task of protein-phenotype co-mention classification.

There are still many avenues to work in this area. We utilized syntactic features extracted from all sentences. However, a potential future work is to incorporate the section titles, e.g., Introduction, Conclusion, etc., to employ only the more informative sentences. The performance of our system is restricted due to the lack of annotated data. Nonetheless, we have access to millions of unlabeled co-mentions through ProPheno. Therefore, we plan to develop a semi-supervised framework that takes advantage of unlabeled data for co-mention classification. Another potential future work is to incorporate larger spans of text, e.g., paragraphs, and to incorporate inter-sentence co-mentions.

## Footnotes

1 https://hpo.jax.org/app

2 http://propheno.cs.montana.edu

3 http://doi.org/10.5281/zenodo.3965127

4 https://github.com/mpourreza/DeepPPPred

5 https://www.uniprot.org

